# Sex-based differences in fibrosis markers in human abdominal aortic aneurysms

**DOI:** 10.64898/2026.10.23.761467

**Authors:** Santiago Elizondo-Benedetto, Ryan Wahidi, Batool Arif, Brigida Rusconi, Maxwell C. Braasch, Dina Ibrahim, Mohamed S. Zaghloul, Omar Soltan, Noah Engles, Ibrahim Kuziez, J. Westley Ohman, Luis A. Sanchez, Mohamed A. Zayed

## Abstract

**Background:** Abdominal aortic aneurysms (AAA) are a complex degenerative vascular disease. The role of fibrosis in AAA progression is unclear. Our study evaluated the impact of fibroblast activation protein (FAP), collagen, and vimentin on AAA fibrosis and progression. We hypothesized that FAP may influence fibrosis and disease severity in AAA, with potential sex-specific differences.

**Study Design:** Multi-centered transcriptomic analysis of aortic samples from 15 human AAA and 5 non-AAA was performed to evaluate cell-specific fibrosis gene expression profiles. Histopathological and molecular analysis of 26 male and female AAA samples (mAAA and fAAA) and 29 male and female normal abdominal aortas (mNAA and fNAA) was performed to assess whole-tissue fibrosis markers and signaling.

**Results:** Single-cell RNA sequencing (scRNA-seq) analysis identified eight major cell clusters with distinct sex-based distributions. Although vascular smooth muscle cells (VSMCs) and fibroblast clusters had reduced counts in both male and female AAAs, they more highly expressed *FAP* and *COL1A1* genes, particularly in females. Whole-tissue analysis demonstrated elevated extracellular matrix deposits with markedly increased collagen type 1 in both mAAA and fAAA. Alpha-smooth muscle actin (α-SMA)-positive VSMCs were also reduced in mAAA and fAAA, but FAP was notably elevated, and robustly in fAAA. FAP-positive fibroblasts in fAAA exhibited increased expression of fibrosis markers, suggesting a unique pro-fibrotic phenotype.

**Conclusion:** Our study reveals a sex-based difference in AAA tissue, with elevated fibrosis markers and increased FAP levels in females, suggesting its significant role in disease progression. These findings underscore the potential of FAP as a unique tissue biomarker for AAAs and warrants further investigation into novel targeted therapies that aim to stabilize AAAs, particularly in female patients.

## Introduction

Abdominal aortic aneurysms (AAA) are a chronic degenerative vascular disease characterized by a progressive dilation of the abdominal aorta that results from an imbalance in extracellular matrix (ECM) synthesis and degradation.^1^ Biological sex is a critical factor in the progression and rupture potential of AAAs.^2,3^ Despite a lower prevalence, females with AAAs experience rupture at smaller diameters compared to their male counterparts, highlighting sex-specific vulnerabilities in AAA pathophysiology.^3–5^ This sex-based difference in rupture risk is reflected in contemporary clinical guidelines of diameter thresholds for surgical intervention,^6,7^ but its molecular and structural underpinning is poorly understood.

For patients with AAAs smaller than the diameter threshold for surgical intervention, no specific therapeutic interventions currently exist to decrease the risk of rupture, and these patients are consequently managed expectantly.^8,9^ Despite advancements in diagnostic imaging, accurately stratifying the risk of AAA rupture remains a significant challenge in the clinical setting.^10,11^ There is a pressing need to identify new molecular and matrix markers that can facilitate both monitoring and therapeutic targeting of AAAs, independent of change in aortic diameter. It is understood that AAA development is marked by the upregulation of matrix metalloproteinases (MMPs), which drive the degradation of critical ECM components, such as elastin and collagen.^12,13^ This inciting pathophysiology perturbs aortic wall structural integrity. Persistent inflammation further exacerbates this weakening, contributing to the progression and potential rupture of AAAs.^11,14^

Fibrosis may play a pivotal role in the pathophysiology of AAA, representing the body’s attempt to repair the aortic tissue following ECM degradation. Activated fibroblasts synthesize collagen and other ECM components, thereby contributing to the fibrotic response.^15^ However, the regenerative potential of fibrosis is often undermined by ongoing inflammation, leading to either further weakening or stiffening of the aortic wall.^16^ Notably, stiffening of the aortic wall is thought to exacerbate aneurysmal dilation and heighten the risk of rupture.^15^ Proteins like fibroblast activation protein (FAP), a serine protease expressed by activated fibroblasts, has been highlighted as a potential crucial regulator of fibrosis in cardiovascular disease.^17–20^ Understanding the complex interplay between FAP, ECM degradation, fibrosis, and inflammation is crucial for developing improved diagnostic markers and effective therapeutic strategies to monitor and manage AAA progression. To address this opportunity, this study aimed to explore the role of fibrosis markers, particularly FAP, Collagen 1, and Vimentin in the remodeling of the ECM in AAAs.

## Methods

### ScRNA-seq Data Analysis

Three publicly available scRNA-seq datasets (GSE166676, GSE237230, GSE226492) comprising AAA and non-AAA human samples were obtained from the Gene Expression Omnibus (GEO) for subsequent analysis.^21,29^ The Seurat package (v.5.2.1) was employed to perform combined gene-barcode matrix analysis of a total of 20 human aortic samples (16 unique and 4 repetitions), a doublet-filtering algorithm (scDblFinder) was applied, and the dataset was integrated using the FastMnn batch correction (**Figure S1A** and **Table S1**).^43^ Cells expressing 200-5000 genes were retained, and those with mitochondrial gene expression exceeding 30% were excluded. Principal component analysis was performed using 12 principal components. Hierarchical clustering and sex-specific gene expression analyses were conducted to differentiate male and female samples, performed as previously described.^24^ Biological sex identification helped identify a total of 11 mAAAs, 4 fAAA and 5 NAA, (**Figure S1B&C**). Clustering analyses were performed to normalize the data, identify variable gene expression, and scale the normalized data. Uniform Manifold Approximation and Projection (UMAP) plots were generated based on selected principal components to visualize the data distribution. Clusters of cells were annotated to corresponding cell types by matching them to cell type-specific gene markers. Statistical significance was determined using Bonferroni-adjusted p-values, with a false discovery rate (FDR) adjusted p-value threshold of <0.05 using Wilcox test.

### Patient Selection

We conducted a retrospective analysis of human aortic tissues obtained from the Washington University vascular tissue biobank. Tissue collection and banking were performed under Washington University Institutional Review Board (IRB) approval #201309043, with informed consent obtained from all patients or their legally authorized representatives. The retrospective analysis reported in this study was additionally reviewed and approved by the Washington University in St. Louis Institutional Review Board under study-specific IRB approval #202609109.

For this study, AAA tissue specimens were obtained from mAAA (N=17) and fAAA (N=9) patients undergoing open AAA repair (**Table 1** and **Table S3**). Control normal abdominal aorta from mNAA (N=14) and fNAA (N=15) subjects were included for comparison. These tissues were collected from abdominal solid organ donors, and their de-identified demographic information was obtained from the U.S. Organ Donation and Transplantation System (UNOS). Furthermore, we collected plasma samples from a subset of the mAAA (N=7) and fAAA (N=4) subjects included in this study, as well as from healthy control individuals. For the healthy volunteers (HV) control group, plasma samples were collected from four human subjects who were not diagnosed with AAA and had undergone thoracic outlet syndrome (TOS) surgery (**Table S4**). This study was conducted in accordance with the ethical principles of the 1964 Declaration of Helsinki and its later amendments or comparable ethical standards.

**Table 1.** Demographics for biobank tissue samples of mNAA, mAAA, fNAA and fAAA.

|  |  | mNAA<br>(N = 14) | mAAA<br>(N = 17) | P value | fNAA<br>(N = 15) | fAAA<br>(N=9) | P value |
| --- | --- | --- | --- | --- | --- | --- | --- |
| **Age** mean (SD) |  | 44.2 (16.5) | 68.1 (6.0) | **<0.001** | 53.5 (14.4) | 67.0 (7.3) | **0.02** |
| **Race** | White | 8 (80%) | 15 (88%) | 0.61 | 9 (90%) | 8 (89%) | 1 |
|  | African American | 2 (20%) | 2 (12%) | 0.61 | 1 (10%) | 0 (0%) | 1 |
| **Smoking Status** | Former | 3 (21%) | 6 (35%) | 0.46 | 2 (13%) | 4 (44%) | 0.15 |
|  | Active | 4 (29%) | 9 (52%) | 0.27 | 4 (27%) | 4 (44%) | 0.41 |
| **Comorbidities** | Hypertension | 6 (43%) | 16 (94%) | **0.004** | 5 (33%) | 7 (78%) | 0.089 |
|  | Diabetes Mellitus | 1 (7%) | 2 (12%) | 1 | 2 (13%) | 0 (0%) | 0.51 |
|  | CAD | 3 (21%) | 4 (24%) | 1 | 0 (0%) | 2 (22%) | 0.13 |
| **Medications** | Statins | 2 (14%) | 15 (88%) | **<0.001** | 1 (7%) | 6 (67%) | **0.004** |
|  | Metformin | 1 (7%) | 7 (41%) | **0.04** | 0 (0%) | 0 (0%) | 1 |
|  | Beta-Blocker | 1 (7%) | 4 (24%) | 0.35 | 2 (13%) | 6 (67%) | **0.02** |
|  | ACE-Inhibitor | 1 (7%) | 4 (24%) | 0.35 | 2 (13%) | 3 (33%) | 0.33 |
|  | Antiplatelets | 1 (7%) | 13 (76%) | **<0.001** | 1 (7%) | 7 (78%) | **<0.001** |
|  | Anticoagulant | 2 (14%) | 1 (6%) | 0.58 | 1 (7%) | 1 (11%) | 1 |
| mNAA = male normal abdominal aorta. mAAA = male Abdominal Aortic Aneurysm. fNAA = female normal abdominal aorta. fAAA = female Abdominal Aortic Aneurysm. SD = Standard Deviation. CAD = coronary artery disease. ACE = Angiotensin-Converting Enzyme (ACE). |  |  |  |  |  |  |  |

### Tissue and Blood Processing

Aortic wall tissue was collected from the operating room as previously reported.^37,44^ Tissue samples were obtained from individuals with non-ruptured AAA, RAAA, or NAA. Specimens were immediately placed in cups filled with cold saline and promptly transferred on ice to the vascular surgery research laboratory for further processing, following established protocols.^24^ Blood samples were collected prior to surgery from individuals undergoing open AAA surgery (mAAA and fAAA) and from those undergoing TOS surgery (HV/TOS; **Table S4**). For plasma preparation, blood was collected into sodium heparin tubes and centrifuged for 15 minutes at 3000 rpm to separate the plasma.

### Histopathology and Immunofluorescence

Tissue sections (5 µm) from mNAA, fNAA, mAAA, and fAAA were stained using MT and imaged with a NanoZoomer HT 2.0 (Hamamatsu Photonics; Shizuoka, Japan). These tissues were subsequently subjected to immunofluorescence staining for various fibrosis markers, including fibroblast activation protein (FAP; LifeSpan Biosciences INC, LS-C413071, 1:150), alpha-smooth muscle actin (Sigma, A2547, 1:500), Vimentin (VIM; Biotechne, MAB21052, 10 µg/ml), and type 1 collagen (COL1; Proteintech, 14695-1-AP, 1:1000). Tissue sections were meticulously deparaffinized, rehydrated, underwent antigen retrieval, and were blocked with 10% serum. Immunostaining was then performed using primary antibodies, followed by Cy3 donkey anti-rabbit and Alexa Fluor 647 donkey anti-mouse secondary antibodies (Jackson ImmunoResearch Laboratories, 1:500). The sections were mounted with Fluoroshield mounting medium containing DAPI (Sigma-Aldrich). All slides were imaged using the THUNDER Imager Tissue Microscope System (Leica Microsystems), ensuring high-quality visualization of the staining results.

### ELISA and Immunoblotting

Protein was extracted from mNAA, fNAA, mAAA, and fAAA tissue using RIPA buffer containing protease inhibitor (Sigma #MCL1). Protein quantification was performed using the Bradford assay. Each protein sample (25 µg) and plasma samples (diluted 1:200) from each group were analyzed using the Human FAP sandwich ELISA kit (RayBiotech, ELH-FAP-1) according to the manufacturer’s instructions. Additionally, 25 µg of protein from each sample was separated via electrophoresis on a 4-12% polyacrylamide gel and then transferred to a PVDF membrane. The membranes were subsequently incubated with antibodies targeting fibroblast activation protein (FAP; LifeSpan Biosciences INC, LS-C413071, 1:150), alpha-smooth muscle actin (Sigma, A2547, 1:500), Vimentin (Biotechne, MAB21052, 10 µg/ml), type 1 collagen (Proteintech, 14695-1-AP, 1:2000), and Vinculin (7F9; Santa Cruz, sc-73614, 1:1000).

### Statistics and Reproducibility

Continuous variables are presented as the mean ± SD, and categorical data as counts and proportions. For comparisons between two groups, a two-tailed Mann-Whitney test was performed. Categorical variables were assessed using Fischer exact tests to evaluate associations. For correlation analyses, Pearson correlation was employed for approximately normally distributed continuous variables. MT staining of cross-sections was analyzed using ImageJ software for color deconvolution, threshold adjustment, and region of interest assessment to quantify collagen deposits, as previously described.^45^ IF intensity and colocalization were determined by two blinded observers with analysis conducted in QPath (v. 0.6.0).^46^ Data was considered statistically significant with p ≤ 0.05. All statistical analyses and graphical data representations were conducted using GraphPad Prism (v.10.6.1, San Diego, CA, USA), RStudio (v.4.3.0, Posit Team, 2025).

## Results

### Human single-cell transcriptomics reveals enhanced pro-fibrotic profiles

A multi-centered transcriptomic analysis was conducted by integrating three publicly available single-cell RNA sequencing (scRNA-seq) datasets.^21–23^ A total of 20 samples was consolidated (**Figure S1A** and **Table S1**). To distinguish male and female samples sex-specific gene expression analysis was performed as previously described (**Figure S1B&C** and **Table S1**).^24^ In total, 87,661 high-quality cells from 5 male normal abdominal aortas (NAA), 11 male AAAs (mAAA) and 4 female AAAs (fAAA) were included in our analysis (**Figure 1A**). Eight major cell clusters were identified: vascular smooth muscle cells (VSMCs), fibroblasts, endothelial cells, macrophages, neutrophils, T cells, B cells and plasma cells, utilizing accepted canonical markers (**Figure 1B** and **Figure S2A).**^16,24–26^ Despite similar gene expression profiles between VSMCs and fibroblasts, we distinguished them in initial clustering through high expression of *ACTA2*, *MYH11* and *CNN1* in VSMCs and low expression in fibroblasts (**Figure S2A&B**). Additionally, fibroblasts exhibited higher expression of *PDGFRA* and *FBLN1* (**Figure S2A&C**). Cell-type distribution analysis demonstrated a marked variability between NAA, mAAA and fAAA groups. While immune cell types demonstrated a relative increase percentage in both mAAA and fAAA groups, structural cells exhibited a notable decrease compared to NAA (**Figure 1C&D**). Interestingly, VSMCs and fibroblasts represented the most striking differences, and both mAAA and fAAA showed a significant decrease in the overall percentage of cell counts (**Figure 1C&D**). Since fibrosis genes *FAP*, *COL1A1, COL1A2* and *VIM*, were predominantly expressed in these two cell types (**Figure 1E**), we focused on a VSMCs/fibroblast-specific subcluster analysis (**Figure 1F**). This revealed that despite the lower cell count, *FAP* was significantly upregulated in both mAAA and fAAA compared to NAA controls (p<0.0001; **Figure 1G**). Notably, female AAAs exhibited significantly higher levels of *FAP, COL1A1* and *COL1A2* compared to male counterparts with AAAs and NAA (p<0.0001), suggesting a more pronounced pro-fibrotic profile in fAAAs (**Figure 1G-I**). In contrast, *VIM* did not show significant differences among the analyzed groups (**Figure 1J**).

**Figure 1.**
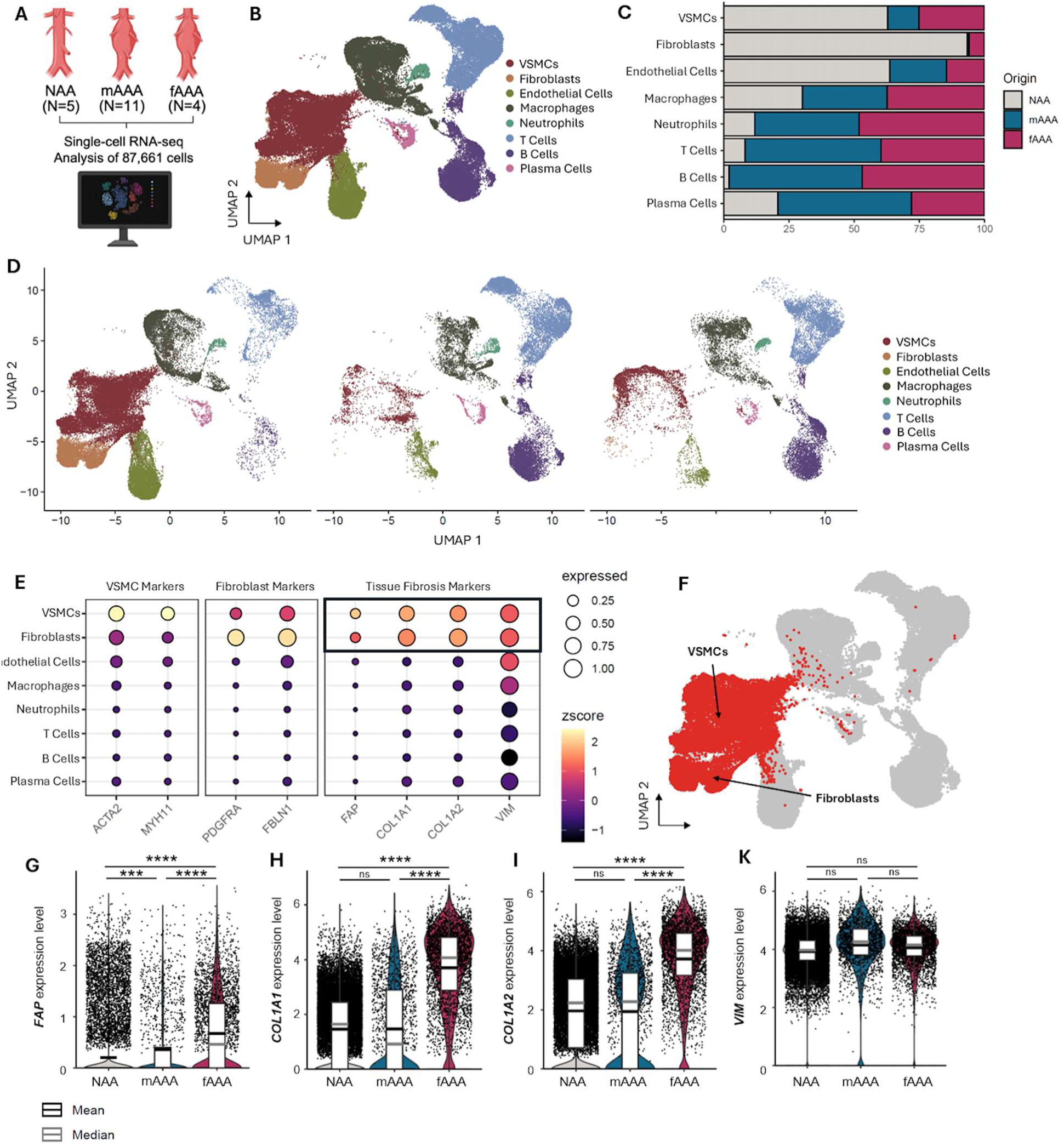
ScRNA-seq analysis of fibrosis markers in AAA tissue and normal aorta. (**A**) 5 normal abdominal aorta (NAA), 11 male AAAs (mAAA) and 4 female AAAs (fAAA) samples were included in the analysis. Panel A was made using Biorender.com (**B**) Uniform manifold approximation and projection (UMAP) plot illustrating identified cell types following unbiased clustering. (**C**) Cell-type distribution analysis represented by the percentage of normalized cell counts, separated by group. (**D**) UMAP plot of identified cell types, separated by group. (**E**) Dot plot of two canonical gene markers for VSMCs and fibroblasts, fibroblast activation protein (*FAP*), collagen type 1A1 (*COL1A1*), collagen type 1A2 (*COL1A2*) and vimentin (*VIM*). (**F**) UMAP of selected subclusters (VSMCs and fibroblasts) for focused analysis. Violin plots of gene expression levels in the VSMCs/fibroblast sub-cluster for (**G**) *FAP*, (**H**) *COL1A1*, (**I**) *COL1A2* and (**J**) *VIM*. Data are presented as mean **±** SEM and median. The Wilcox test was performed, and Bonferroni-adjusted p-values were used to determine significance. Non-significant (ns), *p<0.05, **p<0.01, ***p<0.001, ****p<0.001.

### Demographic and clinical characteristics of human AAA and NAA tissue samples

Human AAA tissue samples (N=26) were collected from our institutional biobank. The male group (mAAA=17) had a mean age of 68.1 ± 6.0 years and the female group (fAAA=9) had a mean age of 67.0 ± 7.3 years (**Table 1**). The majority of participants were white (88%) and had a history of smoking (38% former and 50% active smokers) (**Table S2**). The average diameter of the AAA at the time of collection was 6.09 ± 1.41 cm in mAAA and 5.94 ± 1.13 cm in fAAA (p=ns, **Table S3**). Comorbidities, including hypertension, diabetes, and coronary artery disease, did not differ between male and female AAA patients (**Table S3**). Although beta-blocker use was significantly higher in female AAA patients compared to males (67 vs 24%, p = 0.046), other relevant medications showed no significant differences between groups (**Table S3**).

In contrast, NAA tissue samples (N=29) were collected from organ donors, who were significantly younger compared to AAA patients (49.0 ± 15.9 vs 67.5 ± 6.4 years old, p<0.001; **Table S2**). The male NAA group (mNAA, N=14), had a mean age of 44.2 ± 16.5 years and the female NAA group (fNAA, N=15) had a mean age of 53.5 ± 14.4 years (**Table 1**). Similarly, NAA samples were predominantly from white subjects (85%) and had a slightly lower proportion of smoking history (17% former and 28% active smokers), however the difference was not statistically significant (**Table S2**). Hypertension was significantly less common in NAA subjects compared to AAA patients (38% vs 88%, p<0.001; **Table S2**). Additionally, the use of medications, including statins, metformin, beta-blockers and antiplatelets, was significantly higher in AAA patients when compared to NAA subjects (p<0.001, p = 0.012, p = 0.015 and p<0.001 respectively; **Table S2**). The incidence of other comorbidities and medications were slightly lower in the NAA group compared to the AAA group (**Table S2**).

### Human whole-tissue has increased collagen content in mAAA and fAAA

Histological analysis was performed using Masson’s trichrome (MT), collagen type 1 and vimentin immunofluorescence (IF) staining, and Western blot (WB) on 10 mNAA, 11 fNAA, 12 mAAA and 7 fAAA samples (**Figure 2A**). MT quantification revealed significantly higher collagen deposits in mAAA compared to mNAA (20.7 ± 7.7 vs 3.8 ± 3.8, p<0.0001) and in fAAA compared to fNAA (18.1 ± 4.4 vs 3.5 ± 3.5, p = 0.0001) (**Figure 2B&C**). No significant differences in collagen deposits were observed between male and female AAA samples or between male and female NAA samples (p = ns, **Figure 2B&C**). Analysis of collagen type 1 demonstrated significantly higher deposits in mAAA compared to mNAA using IF staining (3.8 ± 2.1 vs 1.8 ± 1.2, p = 0.04; **Figure 2D&E**) and WB quantification (p = 0.004; **Figure S3A&B**). Similarly, collagen type 1 was significantly higher in fAAA compared to fNAA using IF (4.9 ± 0.7 vs 3.3 ± 1.5, p = 0.02; **Figure 2D&E**) and a slightly higher, though not significant, using WB quantification (**Figure S3A&B).** Although fAAA had the highest collagen type 1 deposits, it was not significantly higher when compared to mAAA counterparts. In contrast, collagen type 1 was significantly higher in fNAA compared to mNAA using both IF (p = 0.0343, **Figure 2D&E**) and WB quantification (p = 0.0028, **Figure S3A&B**). Overall MT, IF and WB analyses demonstrated higher collagen deposits, predominantly driven by collagen type 1, in both mAAA and fAAAs compared to their NAA counterparts; however, these histological and protein comparisons are unadjusted for age. On the other hand, analysis of vimentin through IF and WB quantification, indicated an increased trend in mNAA, but no significant differences were observed (**Figure 2F&G** and **Figure S3A&C**).

**Figure 2.**
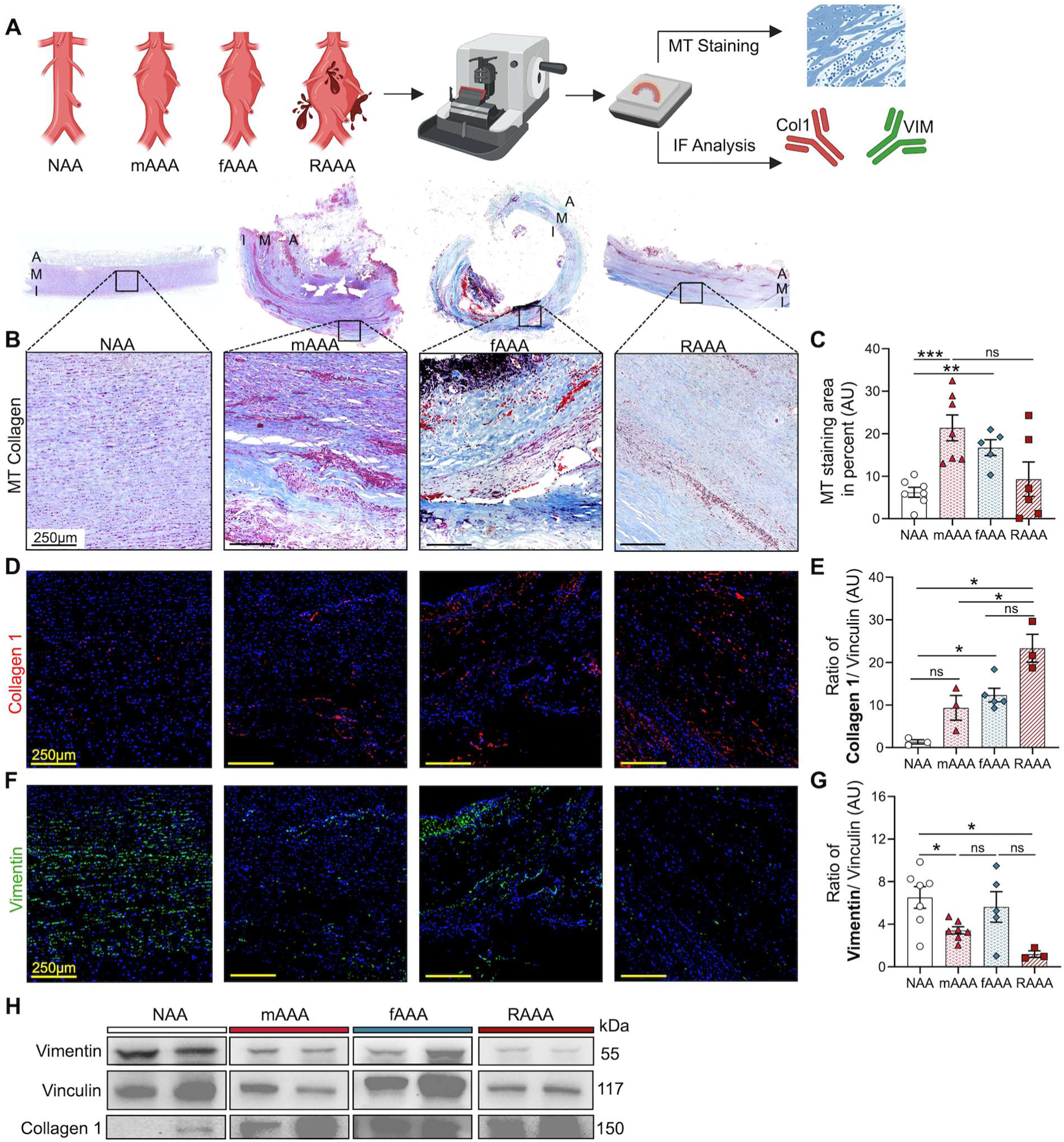
Collagen and vimentin content in human AAAs and NAA. (**A**) Histopathology analysis was performed on mNAA, fNAA, mAAA and fAAA to evaluate for fibrosis markers. This was performed using complementary Masson trichrome (MT), immunofluorescence (IF) staining, and Western blot analysis. Panel A was made using BioRender.com. (**B**) MT staining of human abdominal aortic tissue for all groups with 5x and 40x magnification. Areas with blue staining show regions with higher collagen deposits. (**C**) MT staining quantification via mean intensity divided by the area in percent. (**D**) IF staining and quantification (**E**) of cell counts percentage using collagen type 1 antibody across all groups. (**F**) IF staining and quantification (**G**) of cell counts percentage using vimentin antibody across all groups. AU = Arbitrary Units. Data presented as mean **±** SD. Ns>0.005, *p<0.05, **p<0.01, ***p<0.001 using Mann-Whitney-test.

### Tissue and plasma FAP indicate a pro-fibrotic signal in female AAAs

Human aortic samples were evaluated using IF and WB analysis for the markers alpha-smooth muscle actin (α-SMA) and FAP to evaluate VSMC and fibroblast-specific pro-fibrotic signals. The presence of FAP was further confirmed using ELISA (**Figure 3A**). IF quantification analysis of α-SMA revealed a significant decrease in VSMC counts in both mAAA and fAAA compared to NAA (mAAA versus mNAA, p = 0.008; and fAAA versus fNAA, p < 0.0001) (**Figure 3B&C**). No significant differences in VSMCs counts were observed between mAAA and fAAA samples or between mNAA and fNAA samples (**Figure 3B&C**). WB quantification of α-SMA also indicated a trend towards decreased content within the aortic wall of both mAAA and fAAA compared to NAA tissue (**Figure S3D&E**).

**Figure 3.**
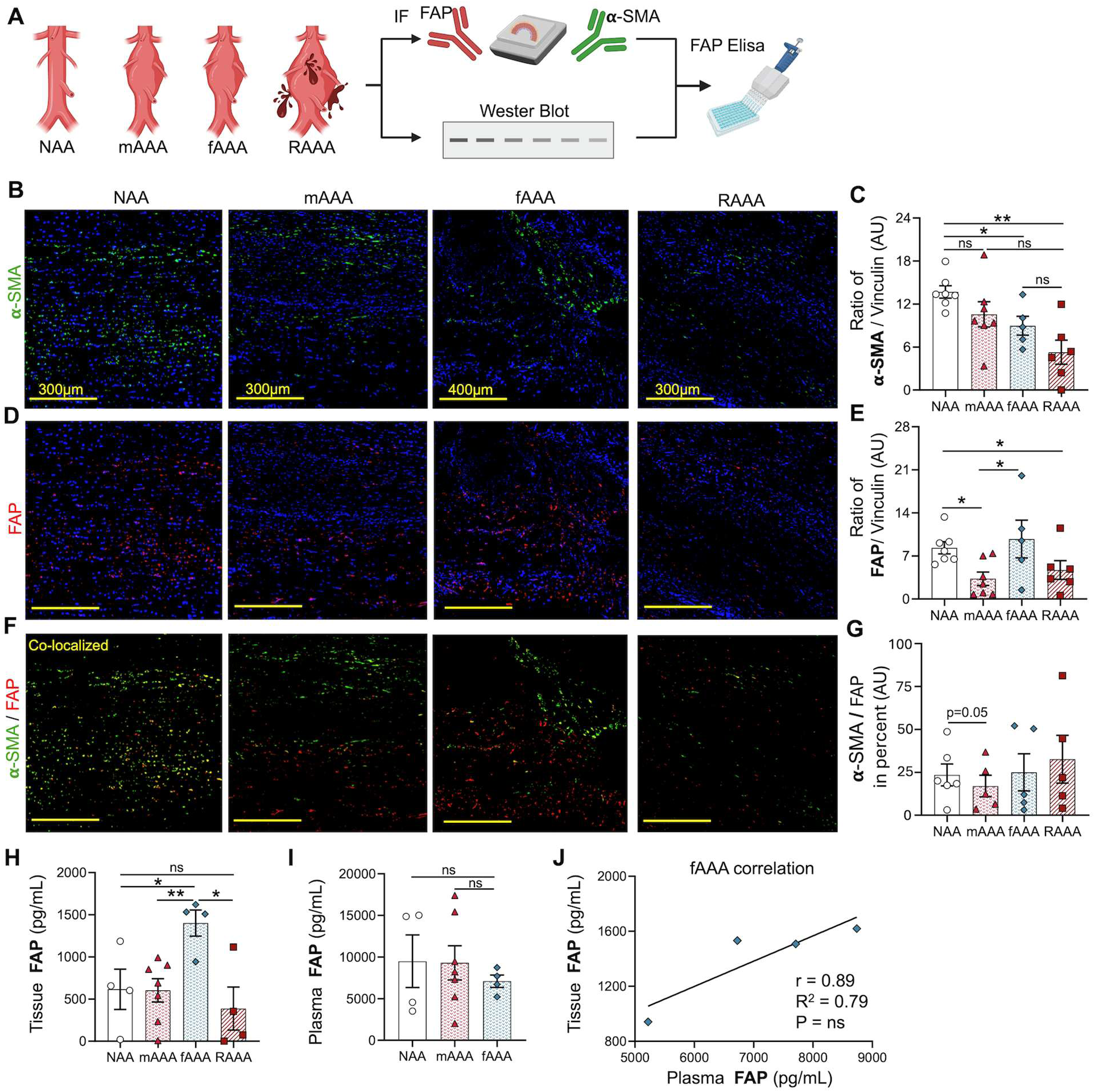
α-SMA and FAP content in human AAAs. (**A**) Histopathology and protein content analysis was performed on tissue from mNAA, fNAA, mAAA and fAAA using immunofluorescence (IF), Western blot and ELISA. Panel A was made using BioRender.com. (**B**) IF staining of human abdominal aortic tissue for all groups with α-SMA (Green), FAP (Red) and DAPI (Blue). Co-localized cells (Yellow) represent VSMCs that express FAP marker. (**C**) IF quantification of VSMC counts using α-SMA antibody across all groups. (**D**) IF quantification of FAP+ cells using FAP antibody across all groups. (**E**) Tissue FAP concentration measured by ELISA. (**F**) Ratio of FAP+ cells that are not VSMCs. AU = Arbitrary Units. Data presented as mean **±** SD. Ns>0.005, *p<0.05, **p<0.01, ***p<0.001 using Mann-Whitney-test.

There was an overall increased quantity of FAP in AAA tissue, particularly in fAAA. Compared to fNAA, FAP levels in fAAA tissue were significantly elevated using IF (39 ± 18 vs. 14 ± 11, p = 0.007; **Figure 3B&D**), WB (10 ± 6 vs. 5 ± 2, p = 0.05; **Figure S3F**), and ELISA quantification (1209 ± 535 vs. 407 ± 213, p = 0.003; **Figure 3E**). In contrast, mAAA tissue showed only a slight increase in FAP via IF (29 ± 17 vs. 18 ± 11; **Figure 3B&D**) and ELISA quantification (567 ± 312 vs. 475 ± 394; **Figure 3E**), with WB revealing a significant decrease (3.3 ± 3 vs. 8.3 ± 3; p<0.05; **Figure S3F**) compared to mNAA. Furthermore, when comparing AAA counterparts, fAAA exhibited a significant increase in FAP levels via WB (p = 0.04) and ELISA (p = 0.004), but only a slight, non-significant increase via IF quantification when compared to mAAA (**Figure 3B, D&E** and **Figure S3F**). This suggested a sex-specific increase in FAP in fAAA tissue.

Although mNAA demonstrated significantly increased FAP via WB (p = 0.003; **Figure S3F**), IF and ELISA quantifications did not demonstrate this (**Figure 3D&E**). Notably, most FAP-positive cells in both mNAA and fNAA were also positive for VSMC, whereas in the AAA tissue, the ratio of FAP-positive cells that were not VSMCs were significantly elevated in both mAAA and fAAA (**Figure 3B&F**). This indicated that the elevated FAP content in AAA, seen robustly in female patients, may be driven largely by fibroblasts rather than VSMCs.

Additionally, plasma FAP levels from mAAA and fAAA were also measured to correlate with tissue levels and were compared to healthy volunteers (HV) (**Table S4**). Plasma FAP levels were unchanged among groups (**Figure S3G**), however, there was a clear positive correlation between tissue and plasma FAP levels in fAAA (r = 0.89), which was not observed in male counterparts (**Figure S3H&I**).

### Activated fibroblasts are prevalent in female AAA tissue

Given the unique distribution of FAP-positive cells in whole AAA tissue analysis, an unbiased analysis of the merged subcluster VSMC/fibroblasts was then performed to evaluate the different cell types that contributed to pro-fibrotic tissue signaling. VSMC/fibroblasts subcluster analysis identified a total of 4 distinct clusters. Two VSMC states were identified: contractile VSMC, predominantly expressing VSMC canonical genes (*MYH11, ACTA2)* while macrophage-like VSMCs expressed inflammatory markers *(CCL3, CCL2* and *HLA-DRA).* Moreover, two distinct fibroblasts (*FBLN2* and *PDGFRA* positive) were also identified as typical fibroblasts, predominantly expressing anti-fibrotic genes (*LAMA2* and *LTBP4*)^27,28^ while activated fibroblast expressed *FAP* and *COL1A2* and other pro-inflammatory genes (*RUNX1* and *CCL2*) (**Figure 4A&B** and **Figure S4**). Activated fibroblasts also expressed decreased *LAMA2* and *LTBP4* relative to the typical fibroblasts, which are genes that are typically reduced in profibrotic states in other tissue types.^27,28^ Interestingly, AAA tissue demonstrated a higher relative abundance of activated fibroblasts compared to healthy aortic tissue (**Figure 4C**), with a paucity of typical fibroblasts. Genes associated with fibrosis (*THY1, RUNX1, THBS1, POSTN, FAP, AEBP1, CLIP2, COMP, COL1A1, MEOX1, and CTHRC1*)^16^ were combined as a module score to compare mAAA and fAAA with normal aortic tissue. Notably higher expression of pro-fibrotic genes was observed in AAA tissue, particularly in fAAAs, in association with the higher proportion of activated fibroblasts (**Figure 4D**). Common pro-fibrotic gene expression was individually compared between the fibroblast subtypes in **Figure 4E**, and this demonstrated elevated expression of each individual gene within activated fibroblasts (p<0.001).

**Figure 4.**
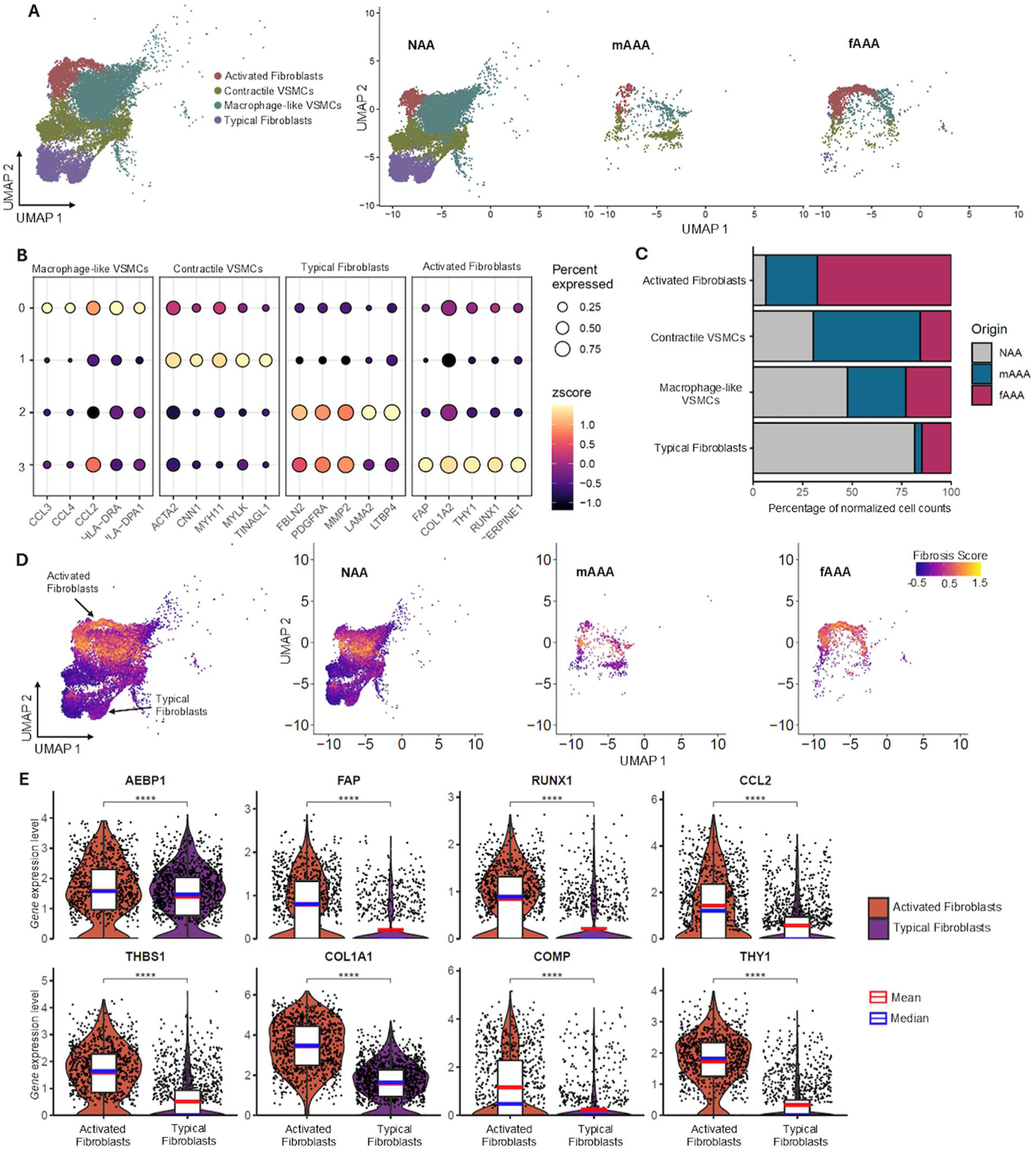
ScRNA-seq analysis of VSMC/Fibroblast sub-clusters in human aortic tissue. (**A**) UMAP plot illustrating the unbiased clustering of the subset containing VSMC and fibroblasts. This revealed 4 cell clusters demonstrated in aggregate and among NAA, mAAA, and fAAA. (**B**) Dot plot of the differentially expressed genes in each cell state: Macrophage-Like VSMC, contractile VSMC, typical fibroblast, and activated fibroblast. (**C**) Relative abundance of each cell state was normalized to the total VSMC + fibroblast content in NAA, mAAA, and fAAA. (**D**) Feature plot of the modulo score of genes commonly associated with fibrosis (*THY1, RUNX1, THBS1, POSTN, FAP, AEBP1, CLIP2, COMP, COL1A1, MEOX1, and CTHRC1*), in aggregate and among NAA, mAAA, and fAAA. (**E**) Violin plots comparing *AEBP1, FAP, RUNX1, CCL2, THBS1, COL1A1, COMP, and THY1* gene expression among activated fibroblasts relative to typical fibroblasts. Data are presented as mean ± SEM and median. The Wilcox test was performed, and Bonferroni-adjusted p-values were used to determine significance, presented as Ns>0.005, *p<0.05, **p<0.01, ***p<0.001.

## Discussion

We conducted a rigorous analysis to elucidate the role of fibrosis markers in human AAA tissue with a particular focus on evaluating potential sex-based differences, revealing several important findings. First, our analysis highlighted the expression of key markers in both male and female AAA tissue: FAP, type I collagen, and vimentin. Second, using a multifaceted approach that leveraged integrated transcriptomics, with subsequent confirmation by protein analysis, histopathology and immunostaining, we identified a significant presence of FAP+ activated fibroblasts in female AAAs. Our findings underscore the potential of FAP as a tissue and serum biomarker for AAAs and warrants further exploration of novel molecular diagnostic strategies and targeted therapeutics that address these observations, particularly in female patients.

ScRNA-seq analysis provided significant insights into the cellular and molecular expression profiles of fibrosis markers in aortic tissue across spectrum of AAA disease severity. Fibroblast-specific clusters exhibited a decreased normalized cell count in both mAAA and fAAA compared to NAA, consistent with prior findings.^29^ Despite the reduced fibroblast population in human AAA samples, gene expression analysis within these clusters revealed elevated levels of fibrosis markers, including *FAP*, *COL1A1* and *COL1A2*. Notably, fAAA demonstrated higher expression of these markers than mAAA (**Figure 1**). These results suggest that these cells demonstrate an increased propensity to express fibrosis markers, despite a lower fibroblast count in AAA tissues. This heightened expression likely represents a compensatory mechanism aimed at mitigating cell loss and promoting the repair of the aortic wall.

Previous studies suggest that the progression and rupture of AAAs is potentially driven by increased collagen turnover. MMPs and other collagenases are highly expressed in AAA tissue, contributing to collagen degradation.^12,13,30^ Historically, literature has indicated an increase in disorganized non-crosslinked collagen within the media of AAA tissues.^31^ To further investigate these findings, we utilized bio-banked human aortic tissue samples from individuals with AAA and healthy aortas, obtained from our institutional vascular biobank – a unique and valuable resource for investigations focused on human vascular pathology. MT staining analysis revealed significantly higher collagen content in both mAAA and fAAA tissue compared to mNAA and fNAA. Consistent with prior literature, we observed no significant differences in overall collagen deposits between mAAA and fAAA groups.^32^ However, collagen type 1 (**Figure 2D&E**) levels were highest in female samples, which correlated with increased gene expression of *COL1A1* and *COL1A2* seen in fibroblasts of fAAAs (**Figure 3H&I** and **Figure 4B&E**). These results highlight dynamic changes in collagen content and emphasize its critical role in maintaining the structural integrity of the aortic wall under pathological conditions. Conversely, we observed slightly decreased vimentin levels in both mAAA and fAAA compared to mNAA and fNAA. Vimentin, a type III intermediate filament protein, is a critical marker for cellular structure, arterial remodeling, and the regulation of fibrosis and wound healing.^33^ This reduction may indicate a diminished capacity for fibroblast migration and activation, which are typically essential for tissue repair and regeneration.

Current literature underscores the significant role of adventitial fibroblasts and their activation in the progression of AAAs.^15^ Adventitial fibroblasts are known to express markers for VSMCs, such as α-SMA. VSMCs also play a crucial role in AAA disease; however, there is ongoing controversy in the literature regarding the identification and distinction of fibroblasts and VSMCs in the aortic wall. Therefore, fibroblasts expressing FAP have recently been identified and found to be elevated in various cardiovascular diseases, as well as in human AAA samples and during AAA progression in various animal models.^19^

In our study, we first identified the presence of VSMCs through IF staining for α-SMA and corroborated these findings with Western blot protein analysis. Similar to vimentin, α-SMA expression was downregulated in mAAA and fAAA compared to mNAA and fNAA, suggesting a reduction in VSMC content in the AAA wall. To specifically assess fibroblasts, we performed FAP staining and protein analysis. Tissue analysis using complementary IF, WB, and ELISA demonstrated consistently upregulated FAP content in female AAAs, while the increase was not significant in mAAA. While recent analysis has highlighted FAP as a crucial marker of AAA progression,^19^ sex-based differences had not previously been demonstrated. Moreover, prior studies lacked robust and comprehensive quantification of FAP levels. One study also reported elevated FAP levels in the serum of patients with AAA compared to those with NAA.^19^ In our analysis, FAP levels in plasma, measured by ELISA, showed no significant differences between NAA and mAAA samples. It is important to note that our study is limited by the small sample size of the blood collections, and that further validation with larger cohorts is necessary to confirm for broader validation in diverse AAA phenotypes.

Interestingly, the co-localization of VSMC and FAP was higher in normal aortic tissue compared to mAAA and fAAA. This finding contrasts with recent studies reporting increased co-localization in the AAA wall of human subjects, despite lacking quantitative analysis.^19^ Our results align with previous reports of decreased α-SMA expression in animal models, as demonstrated with Western Blot.^34,35^ The decrease in co-localization markers suggests potential fibroblast activation and migration, which are crucial for arterial remodeling and tissue repair. Furthermore, our findings highlight that FAP+ fibroblasts tend to localize deeper in the AAA intima and are virtually non-existent in the superficial intimal layer.

Remarkably, this sub-population of FAP+ fibroblasts is consistent with previous findings and shows that many FAP+ cells do not co-localize with α-SMA, indicating that FAP may originate from various cell types including but not limited to VSMCs.^16,20,36^ The role of FAP+ fibroblasts is especially significant as recent studies have established their crucial involvement in cardiac fibrosis following chronic and acute heart failure.^16^ FAP+ fibroblasts are demonstrated to play pathological roles linked to inflammation, via activation by pro-inflammatory macrophages expressing the CC-chemokine receptor type 2 (CCR2).^16^

Our previous investigations clearly link CCR2-mediated inflammation to AAA disease progression.^24,37,38^ Here we further explored how VSMC/fibroblast subpopulation are also influenced by inflammation signaling in AAAs and are associated with disease severity. In our analysis we identified 4 distinct sub-clusters, including a unique activated fibroblast cluster predominantly present in fAAAs (**Figure 4**). Remarkably, the *FAP* gene was significantly upregulated in these activated fibroblasts, suggesting a distinctive phenotype. Applying an established fibrosis score for cardiac fibroblasts, we confirmed this unique phenotype’s presence and upregulation specifically in fAAA tissue.^16,39^ Moreover, these activated *FAP* fibroblasts also exhibited high expression of genes such as *AEBP1*, *RUNX1*, *THBS1*, *COL1A1*, *COMP* and *THY1*. Interestingly, FAP fibroblasts also demonstrated elevated *CCL2* expression, which leads to the recruitment of macrophages via CCR2.^16,40–42^ Overall, the genotype of *FAP* fibroblasts stand outs as unique marker that could potentially improve the identification of highly fibrotic AAAs, particularly in female patients.

Our study highlights the importance of considering sex-specific biomarkers in the ongoing translational investigation of AAA disease pathology – particularly when evaluating the impact of tissue fibrosis on AAAs. Given the stark differences in gene expression and cellular profiles between mAAAs and fAAAs, it will be imperative that future studies include sex-specific stratification of data and explore differences in outcomes between molecular and biochemical endpoints. Larger cohorts of pooled aortic tissue data sets from both males and females are needed to further validate the mechanisms driving the sex-specific roles of FAP in AAA disease progression and fibrosis.

## Limitations of the Study

While FAP shows promise as a tissue marker in AAA, further study is required to understand its role and efficacy as a potential therapeutic target. A central limitation of our study is the lack of functional validation of FAP inhibition in an in vivo animal model to elucidate its role in aneurysm growth, derangement in tissue biomechanical properties, or rupture risk. Likewise, additional studies are required to precisely define upstream signaling leading to persistent fibroblast activation.

Additional limitations of our study include first, the absence of age and sex-matched healthy controls for the scRNA-seq cohort, which limits our ability to draw definitive comparisons and potentially confounds the interpretation of our grouped results. This was compounded by the relatively small sample size of fAAA patients in the scRNA-seq cohort (n=4). However, to address this, we were able to collect bio-banked and donor tissue from both male and female NAA, as well as a robust set of fAAA patients, which corroborated our scRNA-seq findings and demonstrated consistency between the different patient populations that were studied. Importantly, the NAA and AAA cohorts could not be perfectly age-matched, notable given the association between age and tissue fibrosis, and warrants further investigation to highlight the independent impact of AAA mechanobiology on vessel wall fibrosis. The limited sample size of patients for plasma FAP levels limits our capacity to draw definitive conclusions. However, despite this constraint, our study does provide valuable insights that extend prior observations that also evaluated plasma FAP in a small population cohort.^19^ Together, our study provides a novel investigation of tissue fibrosis markers in individuals with AAAs, and provides a unique comparative analysis between normal abdominal aorta, mAAA, and fAAA tissue.

## Conclusions

Our analysis advances our understanding of the content of fibrosis markers in AAAs. Using a multi-faceted approach, we demonstrated dynamic changes in collagen content relative to sex-specific AAA disease. Contrary to existing literature, our analysis revealed a markedly upregulated expression of genes responsible for type 1 collagen 1 in fAAA, while vimentin levels were decreased, indicating potential impairments in fibroblast migration and activation. Moreover, our findings highlighted the presence of a sub-population of FAP+ fibroblasts that have diverse origins and have a unique expression profile that may be actively contributing to AAA disease progression. Notably we also observed elevated levels of FAP in AAA tissue of female patients, suggesting this as a potential tissue biomarker for clinical monitoring and fibrosis severity. These findings underscore the importance of evaluating fibrosis markers in AAA disease pathology, and larger cohort studies are needed to further validate our findings. Translational opportunities may leverage our study findings and facilitate targeting of fibrosis markers for AAA disease stabilization and prevention of rupture.

## Acknowledgment

We acknowledge the vascular tissue biobank at Washington University School of Medicine in St. Louis for providing deidentified aortic tissue. This work was supported by grants from National Institute of Health, National Heart Lung and Blood Institute R01HL153436 (M.A.Z.), R01HL150891 (M.A.Z.), and R01HL153262 (M.A.Z.).

## Author contributions

Conceptualization: S.E.B., B.R., M.S.Z., M.A.Z.; Methodology: S.E.B., R.W., B.A.; Investigation: S.E.B., R.W.; Data Collection: S.E.B., B.A., M.C.B., D.I., M.S.Z., O.S., N.E., I.K., J.W.O., L.A.S.; Supervision: B.R., M.A.Z.; Writing – original draft: S.E.B., R.W., M.A.Z.; Writing – review & editing: S.E.B., R.W., B.R., M.C.B., D.I., J.W.O., L.A.S., M.A.Z.

## Supplemental Information

Figure S1 Figure S2 Figure S3 Figure S4 Table S1 Table S2 Table S3 Table S4

## Data Availability Statement

The datasets and computer code utilized in this study are available from the corresponding author upon reasonable request. Computer code utilized was deposited in GitHub (Zayedlab/FAP_Fibrosis_AAAs). The RNA-seq data sets are publicly available in the Gene Expression Omnibus under accession numbers GSE166676, GSE237230, and GSE226492.

